# Poor attentional control as a sex-specific biomarker to assess vulnerability to nicotine addiction in mice

**DOI:** 10.1101/2022.09.29.510104

**Authors:** Maria-Carmen Medrano, Florence Darlot, Martine Cador, Stephanie Caillé

## Abstract

Every day thousands of people smoke a first cigarette, exposing themselves to the risk of becoming addicts. But this risk is not equal from individual to individual, inviting the hypothesis of potential biomarkers for predicting baseline vulnerability to addiction. One property of nicotine is to increase attentional capacities. However, the role of this pro-cognitive nicotinic effect in initiation of habitual smoking is unknown. Here, we investigated whether the differential effects of nicotine on cognitive performance in mice were predictive of sensitivity to nicotine reward and, if so, whether this characteristic was sex dependent. Naïve populations of male and female mice were first assessed for their attentional performances in the attentional cued-Fixed-Consecutive-Number task (FCNcue) in baseline conditions and after nicotine injections (0.15 and 0.30 mg/kg). Next, all mice underwent nicotine-induced conditioned place preference (CPP) in order to evaluate inter-individual differences in nicotine (0.30 mg/kg) reward sensitivity. Our results showed that innately impulsive males, but not females, benefited from the pro-cognitive effect of nicotine and were also subsequently more sensitive to nicotine reward, indicating increased vulnerability to developing nicotine addiction. Females displayed a completely different behavioural pattern, whereby nicotine reward sensitivity was independent of baseline attentional performances. These results suggest that the pro-cognitive effect of nicotine plays a key role in the development of nicotine addiction in males but not females. Moreover, they signal that the cognitive processes and neurobiology underpinning innate impulsivity may differ significantly between males and females.

## 1. Introduction

Tobacco addiction is one of the most widespread addictions in the world and continues to be one of the leading risk factors for premature deaths globally (WHO, 2021). Every year, 8 million people die because of tobacco use. Yet despite the dangers of smoking being highly publicized, every day tens of thousands of people smoke a first cigarette, opening themselves to the risk of becoming addicted to nicotine, the major psychoactive component in tobacco (1). That risk is not equal from individual to individual; however, we lack a complete understanding of what makes one person more likely to become addicted than another.

Previous studies strongly suggest that women and men differ in the nature of their initial nicotine sensitivity (2). Habitual smoking in men seems to be favored by high impulsivity, novelty seeking, and high sensitivity to nicotine reward, while vulnerability to nicotine addiction in women may be more related to nicotine-associated cue reactivity and regulation of negative mood (3). There is also an abundant literature on the existence of sexual dimorphism in the various processes of nicotine addiction in rodents. This dimorphism has been illustrated by differences in the functioning of the mesolimbic dopamine system (4), in the amount of self-administered nicotine (5), and in reward deficit and somatic signs associated with precipitated nicotine withdrawal in rats (6).

Independently of sex, dysfunctions in inhibitory control and attention may predispose an individual to different levels of nicotine addiction (7). While higher nicotine seeking and intake behaviours have been associated with deficits in cognitive functions such as flexibility, behavioural inhibition, planning, or working memory in both humans and rodents (8), the direction of a causal relationship between attentional deficits and nicotine addiction is less clear. We know that nicotine has pro-cognitive properties, notably improving attention, in rodents and humans (9–11). This pro-cognitive effect of nicotine plays a reinforcing role in tobacco addiction and relapse (12). However, little is known about the link between the pro-cognitive effect of nicotine and the initiation of smoking in healthy individuals, rodents or humans.

We previously investigated the predictive relationship between attentional performances and the motivation for nicotine using our established cued-Fixed Consecutive Number task (FCNcue, (13)) and intravenous nicotine self-administration schedules (14) in male rats. Thus, we demonstrated that male rats who pay more attention to reward-predictive cues showed also higher sensitivity to both reinforcing and incentive properties of nicotine self-administration (14). However, the impact of the pro-cognitive effect of nicotine in this higher sensitivity to the drug reinforcing effects and motivation to self-administration, remains unknown. In the present study, we aimed to stablish behavioural biomarkers, related to the innate and drug-altered attentional control of male and female individuals, which will help us to fill the knowledge gap in our understanding of the predisposing factors involved in the vulnerability to the development of nicotine addiction.

The objective of this study was to examine potential sexual dimorphism in the relationship between innate cognitive profiles, the pro-cognitive effects of nicotine and nicotine reward sensitivity. For this, naïve populations of male and female mice were first assessed for their attentional and inhibitory control performances in the FCNcue task in basal as well as in nicotine dose-response conditions. Next, all mice underwent nicotine-induced conditioned place preference (CPP) in order to evaluate inter-individual differences to nicotine reward. Based on this multidimensional analysis of distinct task behaviours to explore nicotine’s attentional and rewarding properties in mice, we were able to identify innate impulsivity phenotypes as a sex-dependent biomarker of vulnerability to develop nicotine addiction.

## 2. Materials and Methods

### 2.1. Animals

A total of 67 C57BL/6 mice, 30 males and 37 females, all 6-8 weeks old at the beginning of the study, were used in this work. Mice were housed in collective cages of 6-9 animals each, on an inversed 12/12 light/dark cycle. Behavioural experiments were performed during the dark phase. Mice were maintained in a controlled-temperature room (22 ± 1 °C) with other cages containing other mice, males and females.

Estrous cycle phase was systematically checked in all females during the FCNcue test phase following the protocol described by Byers and cols; (15). Females were tested across the different stages of the cycles.

All procedures were conducted in accordance with the local ethics committee of the University of Bordeaux and the European Committee Council Directive (approval number 2016012716035720). All efforts were made to minimize animal suffering and reduce the number of animals used.

### 2.2. Apparatus

The Cued-Fixed Consecutive Number (FCNcue) procedure was conducted in 12 identical sound-attenuated and ventilated operant chambers (24 × 16 × 21 cm; Imetronic, France) located in a dimly lit room with a background white noise. Each operant chamber was equipped with two nose-poke holes (NP), one on the right wall and one on the left wall, and cue lights situated above each NP. The floor consisted of stainless-steel bars separated by 0.4 cm. A liquid dipper (licker) was situated on the left wall, next to the NP. 75μl of diluted condensed milk (1:5) solution was delivered in the licker as a reward (See Fig 1 A). The operant chambers were interfaced with a PC computer for data acquisition and storage. The reward delivery, presentation of visual stimuli, and recording of behavioural data were controlled and recorded by POLY software (Imetronic, France).

**Figure 1.**
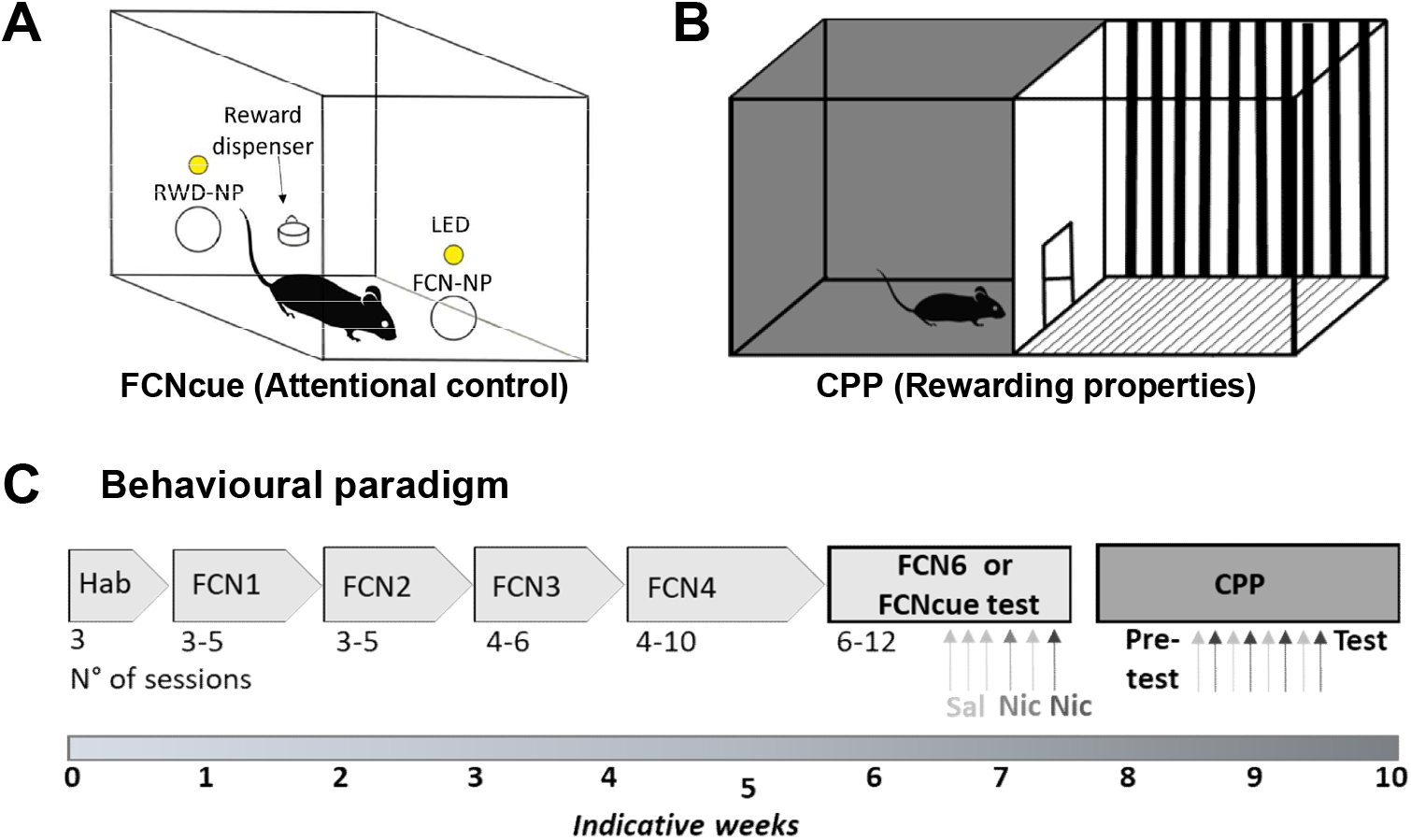
Schematic representation of the behavioural paradigms. A) Schematic representation of the operant cages (Imetroinc ®) used for the Cued-Fixed Consecutive number (FCNcue). B) Schematic representation of the boxes used for the Conditioned Place Preference (CPP) for nicotine. C) Timeline of one experiment. Mice were trained and tested in the FCNcue. In the FCNcue test phase (FCN6cue) mice were injected first with saline (control, light grey arrows) and after with nicotine (0.15 and 0.30 mg/kg, grey and dark grey arrows respectively). Following the FCNcue test, mice underwent the CPP for nicotine in which alternated injections of saline and nicotine 0.30 mg/kg were performed during 8 consecutive days between the pre-test and the test day.

### 2.3. Cued-Fixed Consecutive Number (FCNcue) procedure

The FCNcue schedule measured the ability of mice to carry out a chain of sequential acts (NP) to obtain a reward (13,14). The task required a fixed minimum number of responses (1 for FCN1, 2 for FCN2 and so on to 6 for FCN6) on the NP on the right (from now on, FCN-NP) followed by a response on the second NP, on the left (reward NP, RWD-NP), resulting in one reward delivery (1 optimal chain = 1 reward) (Fig 1 A). The completion of the required number of FCN-NP was signaled by the switching off of the cue light above the FCN-NP. Then, the RWD-NP became active, which was indicated by the switching on of the cue light above it. After nose poking in the RWD-NP, the cue light turned off and the reward was distributed in the licker. Once all the reward was consumed (45 licks required), the cue light above FCN-NP turned on indicating the beginning of a new chain. Premature responding on the RWD-NP prior to the completion of the chain of visits to the FCN-NP, reflected poor inhibitory control and this reset the chain without reward distribution. High rate of optimal chains reflected selective attention to the cue light.

#### 2.3.1. FCNcue training and test

The full experimental timeline is showed in Figure 1 C. The habituation phase consisted in three sessions across three consecutive days. The first day, mice were habituated to empty operant cages (no NP, cue lights or licker) for 30 minutes. The second and third day, mice were habituated to the FCN-NP, the cue light and the reward delivery. Each visit to the FCN-NP resulted in the delivery of a reward. Then, mice underwent FCNcue training during which both, FCN-NP and RWD-NP, were continuously available throughout 1-hour sessions. First, mice had to nose-poke once in the FCN-NP and then once in the RWD-NP to obtain condensed milk (FCN1). After 3 sessions, mice were moved to the next level, FCN2 according to the same criterion. Then to FCN3. After three consecutive stable sessions in FCN3, they moved to FCN4. Finally, mice reached the test condition, FCN6. After the establishment of stable performance in the FCN6, mice were injected with saline (s.c.) for three consecutive sessions as a control In the following sessions, mice were injected with nicotine 0.15 and 0.30 mg/kg, respectively. One session with saline injection was made between the two nicotine sessions (see Fig 1 C). The doses of nicotine were chosen based on previous studies in the laboratory (16). All training and test sessions lasted 1 hour and animals underwent one session per day.

#### 2.3.2. Behavioural measurements

All behavioural data showed in this study were calculated in the FCN6cue, corresponding to the test phase of the task (see Fig 1 C).

The mean percentage of optimal responses [(number of chains of 6 FCN-NP/ (total number of chains of NP) × 100] was an index of attentional performance. Therefore, high percentages of optimal responses denoted good attentional control, whereas low percentage of optimal responses denoted poor attentional control.. The mean percentage of premature responses [(number of chains of 1 FCN-NP) / (total number of chains of NP) × 100] was considered as an index of poor inhibitory control, i.e. impulsivity. Chains of 1 (and not chains of 2, 3 or 4) were chosen to represent premature responses since they are the shortest chains of nose-pokes possible and therefore, which best represent impulsive behaviour in the FCNcue task. Response efficiency was defined as the total number of rewards (i.e. RWD-NP correct) divided by the number of RWD-NP, [(number of rewards) / (total number of RWD-NP) × 100]. High percentages of optimal responses and response efficiency both denote good attentional control since they directly depend on the capacity to perceive the cue light switch off as an indicator of the next action to execute, i.e., selective attention to the reward-predictive cue. Response rate, calculated as the total number of nose pokes per minute [(FCN-NP + RWD-NP) / 60] and the total number of chains of NP, were used as an index of motivation. The activity (total counts) and number of crosses across the chamber were used to measure general locomotor activity. The mean latency to collect the reward was considered a measurement of reward seeking and the sensitivity to the reward. Therefore, the FCNcue test allowed us to obtain behavioural parameters related to attentional control, that we could use as behavioural biomarkers to study vulnerability to nicotine addiction.

### 2.4. Conditioning Place Preference (CPP)

We then performed the CPP for nicotine as previously described (16). An unbiased design was used with two chambered conditioning place preference (CPP) boxes (70 × 80 × 35 cm). The boxes consisted of two side chambers that featured different wall and flooring patterns to make them easily distinguishable (Fig 1B). A 10 × 15 cm door provided access between the boxes. First, mice were submitted to a 15-minute, drug-free, preconditioning test (pretest) in which mice had free access to the two chambers. In a second phase (conditioning), the door between the two chambers was closed. Mice received subcutaneous (s.c.) injection of saline (5 ml/kg) on days 1, 3, 5, and 7 prior to being confined to their vehicle-paired chamber for 30 min. On days 2, 4, 6, and 8, mice received nicotine solution (0.30 mg/kg) immediately prior to confinement in the nicotine-paired chamber for 30 min. Lastly, on day 9, the test session was conducted exactly as in the pretest condition, with mice having free access to the two chambers for 15 min. Animals were tested drug-free (Fig 1 C).

The time spent in each chamber during the pretest and test sessions was counted. A preference score was calculated as the difference between the time spent in the nicotine-associated chamber during the test and the time spent in the same chamber during the pretest.

### 2.5. Drugs

Acute nicotine injections were performed in the test phase (FCN6) of the FCNcue task and every other day in the CPP. Nicotine hydrogen tartrate (Sigma-Aldrich®, France) was dissolved in saline and pH regulated at 7.4. NaOH (Sigma-Aldrich®, France) was used to adjust the pH. Nicotine dose was expressed as nicotine base and we used the following concentrations: 0.15 and 0.30 mg/kg. Nicotine or its vehicle, saline, were injected s.c. 5 min before the session. Mecamylamine (Sigma-Aldrich®, France) was dissolved in saline and injected intraperitoneally (i.p.) at a volume of 5 ml/kg, 5 min before nicotine injection. Amphetamine 0.30 mg/kg (d-amphetamine sulfate, Cooperative Pharmaceutique Française, France) was dissolved in saline and injected i.p. at a volume of 5 ml/kg.

### 2.6. Statistics

The statistical analysis for FCNcue and CPP data was performed with GraphPad software (Prism 9 for Windows, GraphPad Software, Inc., San Diego, CA, USA). For the analysis of data from FCNcue, two-way analysis of variance (ANOVA), followed by multiple comparisons using the Bonferroni post hoc test was used to compare two behavioural parameters between males and females. Repeated measures one-way ANOVA was used to compare within-subject factors (saline versus drug). Upon significant interaction, post hoc analyses were performed using Bonferroni’s post hoc test. Paired t test was used to compare the means of two parameters in the same group of subjects and Unpaired t test to compare two parameters between males and females. In order to identify attentional control biomarkers that are associated with greater vulnerability to developing nicotine addiction, we split male and female groups in three sub-populations based on the mean optimal responses during the control sessions in FCNcue test (FCN6cue). Thus, mice were classified as impulsive (lower thirtieth percentile), medium (medium thirtieth percentile) or good performers (upper thirtieth percentile). For the correlations, we computed Pearson correlation coefficients. One sample t test was used to determine if the preference for nicotine in impulsive and good performers sub-populations and if the simple linear regression showed in correlations were different from “0”.

For the CPP data, Wilcoxon test compared to “0” (chance level) was used to compare the time spent in the nicotine-paired chamber before and after nicotine conditioning. Fisher’s exact test was used to compare the proportion of males and females that show preference for nicotine.

Data are given as mean ± standard error of the mean (s.e.m.). A significance level of p < 0.05 was used for all statistical analyses.

## 3. Results

### 3.1. FCNcue schedule reveals differences in attention and inhibitory control between males and females

First, we addressed the inter-individual variability among males and females regarding their performance in the FCNcue test (or FCNcue6). Performance among individuals, males and females, was heterogeneous: all mice behaved differently, with more or less chains of 6 consecutive FCN-NP (from now on optimal responses) or premature (chains of 1 FCN-NP) responses (Fig 2 A-B). The percentage of optimal responses in the FCNcue test provides information on attentional capacities i.e. how much the animal pays attention to the cue light to perform a chain of actions. Some mice showed a high proportion of optimal responses while others showed a low proportion. Indeed, optimal responses ranged from 12% to 71% in males and 10% to 66% in females. When we compared the performance of females and males, a two-way ANOVA analysis showed a gender x performance interaction (F (1, 46) = 7.482, p = 0.0088) and an effect of performance F (1, 46) = 7.676, p = 0.0080 (Fig 2 C). Males showed a significantly higher proportion of optimal responses than females (40.30 ± 3.9 % vs. 30.0 ± 2.3 %, respectively, p= 0.0427, Bonferroni’s post hoc test), suggesting that males show better attentional performance than females in the FCNcue test. Moreover, when we compared the rate of premature responses in the FCNcue test between males and females we found that females showed a significantly higher proportion of premature responses than males (29.9 ± 2.7 vs. 17.9 ± 2.6 %, respectively; p= 0.0148, Bonferroni’s post hoc test, Fig 2 C).

**Figure 2.**
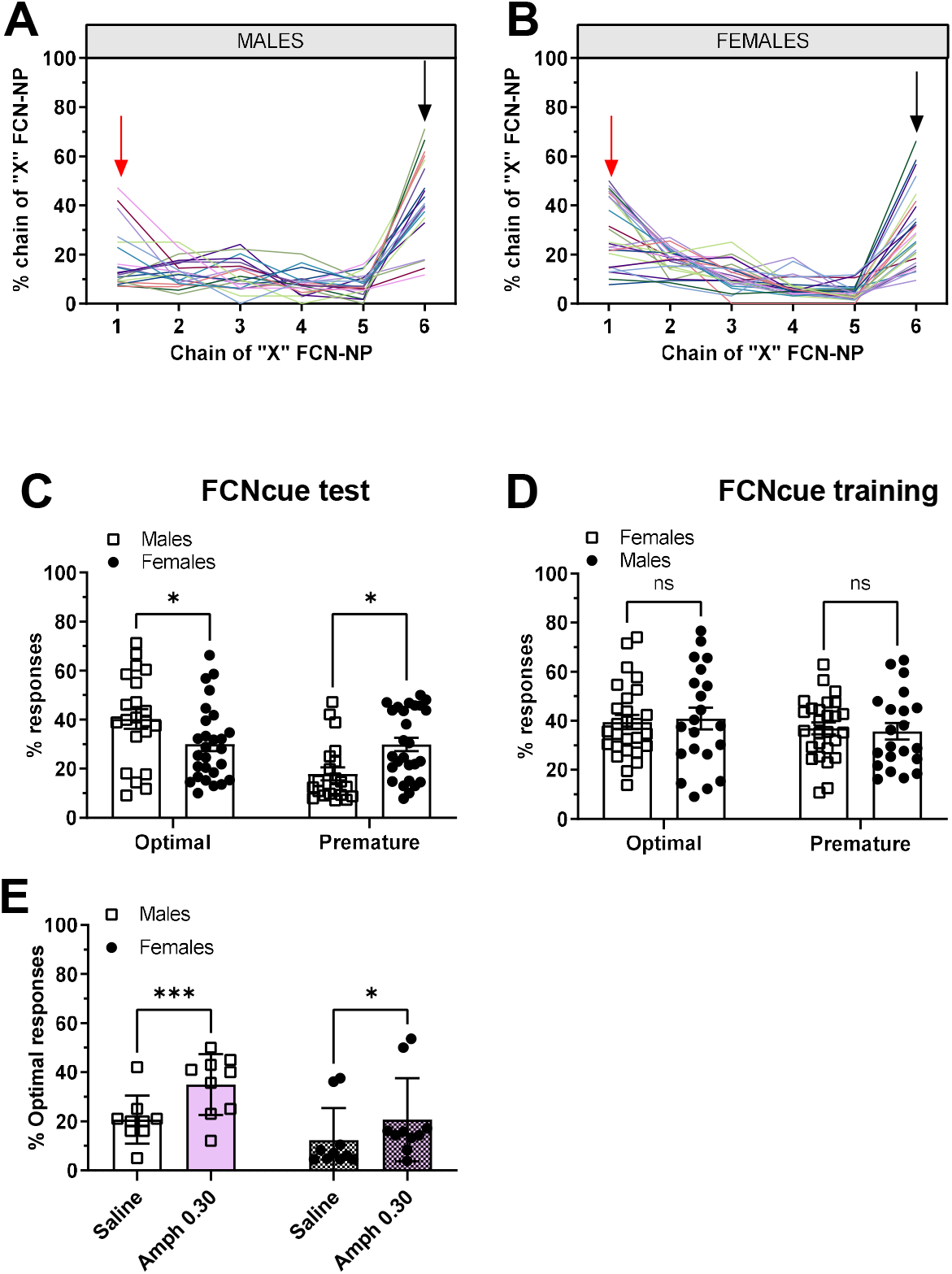
FCNcue schedule reveals differences in attention and inhibitory control between male and female mice. A) Representation of individual performances in a single control session (after saline injection) of FCNcue test (FCN6). Note the difference among individuals in the chain of 1 (red arrow, impulsive responses) and the chain of 6 (black arrow, optimal responses). B) As a group, male performances under the FCNcue schedule are more optimal while females are more impulsive. C) Differences in optimal and impulsive responses between males and females are exclusive of FCNcue test phase, there was no significant differences in the FCNcue training phase. D-E) A single dose of amphetamine (known pro-cognitive agent) improves performances in males and females, validating the FCNcue as a model for studying pro-cognitive effect of drugs in mice. Bars and errors represent mean ± s.e.m. *p<0,05, *** p < 0.005, Bonferroni’s post hoc test.

These results indicate that females, as a group, show poorer attentional control, i.e. impulsivity, than males on the FCNcue schedule. These differences in attention and impulsive behaviours between sexes appear to be specific to the FCNcue test phase (FCN6cue) as there was no difference between sexes performances during the training phase i.e. FCN3cue, mid-term phase of the FCNcue protocol; gender x performance interaction F (1, 46) = 0.08616, p = 0.7704; performance F (1, 46) = 0.7542, p = 0.3897). This indicates that the females’ poor attentional control in the FCNcue test is not related to a learning issue.

In addition to the optimal and premature chains of action, the FCNcue schedule allowed us to extract other parameters related to attentional control, mice locomotor activity, motivation in the task and the incentive value of the reward (see Materials and Methods). We compared the results for these parameters between males and females in control situation (mean of the saline sessions before nicotine injections; Table 1). Males and females showed significant differences in the efficiency of their responses and the number of ineffective RWD-NP. These differences suggested that males paid more attention to the cue whereas females nose-poked more but in a more inefficient way. Females showed higher locomotor activity than males, which could favour their impulsive behaviour. However, the total number of chains of NP and the latency to retrieve the reward were not significantly different between males and females. Altogether, these results indicate that the sex differences observed in the FCNcue task are not due to sex discrepancies in motivation or reward sensitivity, but to differences in attention and inhibitory control.

**Table 1.** Parameters obtained from the FCNcue test for males and females in control conditions. The FCNcue test allowed us to obtain behavioural parameters other than optimal and premature chains, related to attentional performance. Note that males showed to be more efficient than females in the FCNcue test in control conditions, whereas females behaved in a more impulsive way (more RWD-NP). Females also showed a higher general activity during the test (* < 0.05, *** p < 0.005, Unpaired t test).

|  | Males |  |  | Females |  |  | P value |
| --- | --- | --- | --- | --- | --- | --- | --- |
|  | mean | SEM | N | mean | SEM | N |  |
| Empty lick (Nb) | 1044,00 | 111,00 | 21 | 1057,00 | 93,00 | 27 | 0,9409 |
| Total licks (Nb) | 1348,06 | 197,50 | 21 | 1410,42 | 137,58 | 27 | 0,7903 |
| NP-FCN (Nb) | 116,04 | 13,85 | 21 | 114,13 | 14,63 | 27 | 0,9273 |
| **RWD-NP ineffect (Nb)** | **58,48** | 9,57 | 21 | **80,81** | 5,97 | 27 | **0,043497\*** |
| Response rate | 3,10 | 0,37 | 21 | 3,40 | 0,30 | 27 | 0,1273 |
| **Efficiency** | **16,86** | 2,24 | 21 | **10,41** | 1,31 | 27 | **0,011724\*** |
| Nb total chains | 23,48 | 2,37 | 21 | 29,06 | 2,51 | 27 | 0,1245 |
| **Total Activity (Nb)** | **1049,00** | 74,00 | 21 | **1459,00** | 77,00 | 27 | **0,0006\*\*\*** |
| Cage crossings (Nb) | 214,78 | 17,98 | 21 | 210,05 | 14,95 | 27 | 0,8400 |
| Latency (ms) | 3759,00 | 1519,00 | 21 | 5752,00 | 2690,00 | 27 | 0,5531 |

Studies from our laboratory and others have demonstrated that an acute injection of a low dose of amphetamine improves performance on attentional tasks in healthy humans and rodents (13,17). In order to corroborate pharmacologically the FCNcue as a test to measure attentional capacities in mice, we injected a different cohort of male (n = 9) and female (n = 10) mice with amphetamine (0.30 mg/kg i.p.) 5 minutes before the beginning of the FCNcue-test. This dose of amphetamine, 0.30 mg/kg, has shown to be effective to increase attention in mice (17). We found no significant interaction between gender x treatment F (1, 17) = 2.395, p = 0.1402; however, there was an effect of treatment (F (1, 17) = 26.55, p < 0.0001). Bonferroni’s post hoc test showed that amphetamine increased significantly the proportion of optimal responses in males and females (p = 0.0005 and p = 0.0359, respectively; Fig 2 E).

All together, these results show that the FCNcue test can be used to measure attentional control parameters in mice. Moreover, our results show that FCNcue schedule unmasks a sex dichotomy in attention and impulsivity.

### 3.2. Nicotine improves attention and reduces impulsivity in males and females

The continuum of percentages of optimal responses in male and female populations shows the high interindividual variability among drug-free mice. Moreover, previous results also showed significant differences in attentional performances between males and females. Next, we aimed to study if nicotine had an effect on attentional performance and if this effect was similar between the group of males and females. For this, after three control (saline, s.c.) sessions, mice performed the FCNcue test immediately after nicotine injections (0.15 and 0.30 mg/kg, s.c.) and the proportion of optimal and premature responses was analyzed. For clarity’s sake all data related to nicotine effects on attentional parameters in males will be presented first followed by the results in females.

A single injection of nicotine was enough to increase the males’ optimal responses in the FCNcue test (F (2, 40) = 10, 60, p = 0.0002). This pro-cognitive effect of nicotine was due to the dose 0.15 mg/kg (p = 0.0002, Bonferroni’s post hoc test, Fig 3A1), whereas nicotine 0.30 mg/kg did not show a significant effect (p = 0.13, Bonferroni’s post hoc test). Moreover, Nicotine 0.15 mg/kg, but not nicotine 0.30 mg/kg, reduced significantly the amount of premature responses from 17.9 ± 2.6% to 11.2 ± 2.0 % (F (2, 40) = 3, 557, p = 0.044; Nic 0.15 p = 0.0081, Nic 0.30 p = 0.70, Bonferroni’s post hoc test, Fig 3A2). Finally, in order to demonstrate that the increase in attention observed after the injection of nicotine 0.15 mg/kg was due to its action on the nACh receptors, some males received the nACh antagonist mecamylamine (1.5 mg/kg, i.p.) 5 minutes before nicotine injection. Figure 3 A3 shows that Mecamylamine prevented the pro-cognitive effect of nicotine (F (1, 5) = 16. 36, p= 0.0076; Nic 0.15 vs. Meca 1.5 + Nic 0.15 p = 0.0149, Bonferroni’s post hoc test).

**Figure 3.**
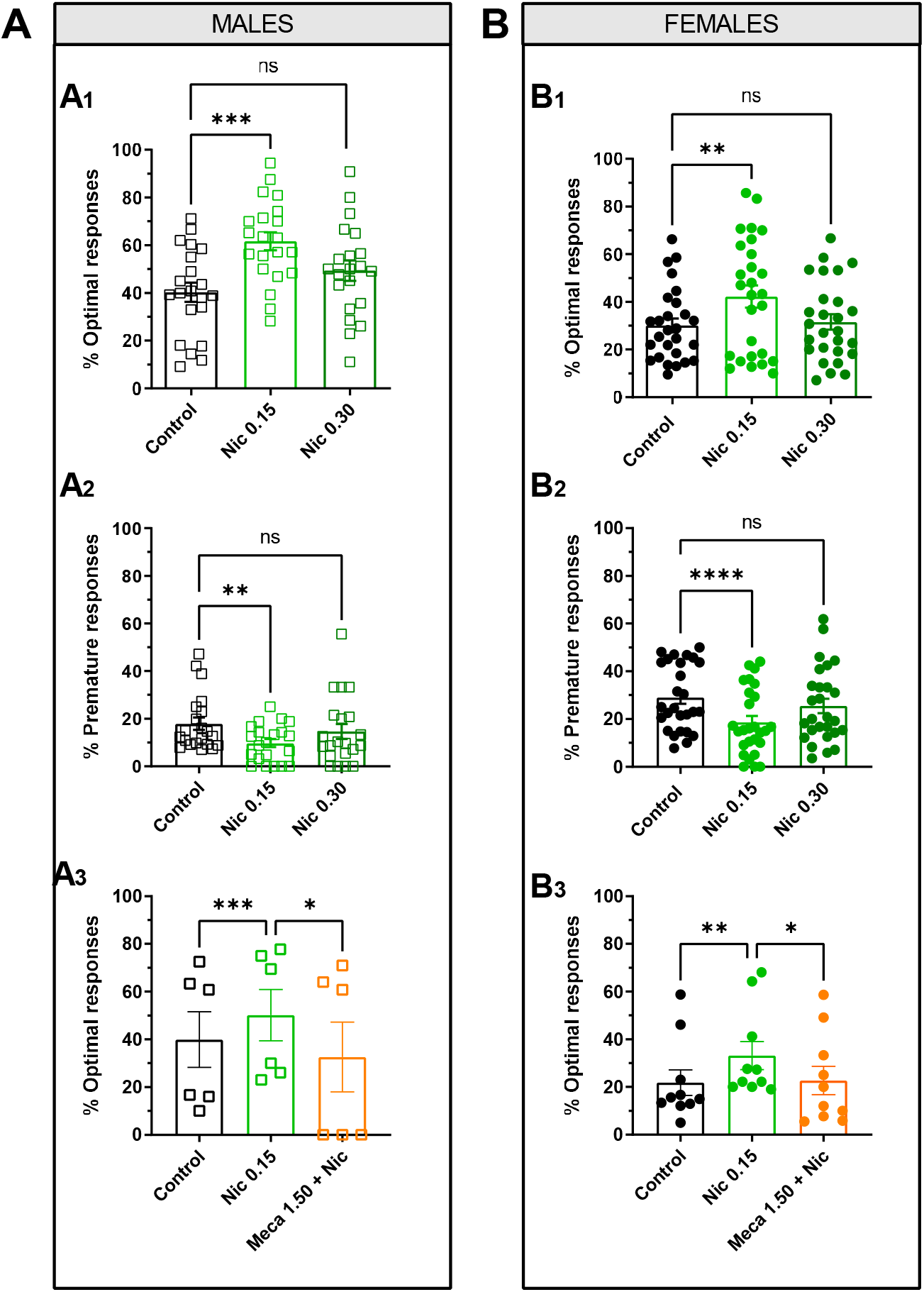
Nicotine improves attention and reduces impulsivity in males and females. A) Nicotine’s pro-cognitive effect in males. A low dose of nicotine (0,15 mg/kg, but not nicotine 0,30 mg/kg) enhances the rate of optimal responses, i.e. improves attention (A1) and reduces the premature ones, i.e. reduces impulsivity (A2) in the group of male mice. Mecamylamine (1.5 mg/kg, nicotinic receptor antagonist) blocks nicotine pro-cognitive effect in the FCNcue task (A3). **B)** Nicotine’s pro-cognitive effect in females. A low dose of nicotine (0,15 mg/kg, but not nicotine 0,30 mg/kg) improves attention (B1) (increase of optimal responses) and reduces impulsivity (B2) (decrease of premature responses) in the group of female mice. Mecamylamine (1.5 mg/kg nicotinic receptor antagonist) blocks nicotine pro-cognitive effect in the FCNcue task. Bars and errors represent mean ± s.e.m. * p < 0.05, ** p < <0.01, *** p < 0.005, **** p < 0.0001, Bonferroni’s post hoc test.

As we described previously, high response efficiency also indicates good attentional control (see Material and Methods). A single injection of nicotine improved males efficiency (F (2, 40 = 3.747, p = 0.0343). This effect was due to nicotine 0.15 mg/kg, whereas the higher dose 0.30 mg/kg showed no significant effect (p = 0.0375 and p = 0.0862, respectively, Bonferroni’s post hoc test; Suppl fig 1 A1). Moreover, the FCNcue test allowed us to extract other parameters that could interfere with attentional control such as general activity, motivation and reward seeking. A single nicotine injection decreased the general activity of males in the FCNcue test (F (2, 40) = 7.571, p = 0.0038). This effect was not due to nicotine 0.15 mg/kg but to the higher dose, 0.30 mg/kg (p = 0.1822 and p = 0.0004, respectively, Bonferroni’s post hoc test, Suppl fig 1 A2). The total number of chains of actions (chains from 1 to 6 NP) is a parameter that measures the motivation of the subject to work for the reward (13). We compared the total number of chains of actions (chains of NP) during the FCNcue test in control conditions and after nicotine injection. The results showed that nicotine had no effect on the total number of chains in males (F (2, 40) = 1.537, p = 0.2299; Suppl fig 1 A3), suggesting that the drug does not change the motivation for the reward. Moreover, the fact that nicotine 0.30 mg/kg shows no effect on the number of chains of actions suggest that the decrease in general activity does not affect their motivation to work on the task. Finally, we also analysed if nicotine had an impact on the latency to collect the reward (condensed milk) once a correct chain of NP was achieved. There was no significant effect of nicotine on the latency, suggesting that the drug has no effect on the reward seeking and the sensitivity to the reward in males (F (2, 40) = 0.6332, p = 0.5302; Suppl fig 1 A4 and B4).

A single dose of nicotine also improved females’ performance under the FCNcue test (F (2, 52) = 12.46, p =0.0002). This pro-cognitive effect was due to nicotine 0.15 mg/kg, p = 0.002, Bonferroni’s post hoc test, Fig 3B1), whereas a higher dose of nicotine, 0.30 mg/kg, did not induce a significant effect (p= 0.9184, Bonferroni’s post hoc test). Nicotine also reduced the percentage of premature responses in female mice (F (2, 52) = 14.33 p < 0.0001). This effect was due to nicotine 0.15 mg/kg, but not nicotine 0.30 mg/kg (p < 0.0001 and p = 0.1081, respectively, Bonferroni’s post hoc test, Fig 3B2). An injection of mecamylamine 5 minutes before than nicotine 0.15 mg/kg injection, prevented the pro-cognitive effect of nicotine in females (F (2, 52) = 9.883, p= 0.0042; Nic 0.15 vs. Meca 1.5 + Nic 0.15 p = 0.0341, Bonferroni’s post hoc test; Fig 3 B3). These results show that nicotine improves attention and inhibitory control in the general population of females by activating nAChRs.

In addition, a single injection of nicotine improved females efficiency females (F (2, 52) = 15.17, p <0.0001). This effect was due to nicotine 0.15 mg/kg, but not the higher dose 0.30 mg/kg (p < 0.0001 and p > 0.9999, respectively, Bonferroni’s post hoc test, Suppl fig 1 B1). Nicotine reduced the general activity in females during the FCNcue test in a dose-dependent manner (F (2, 52) = 20.07, p <0.0001, Control vs. Nic 0.15 p = 0.0027, control vs. Nic 0.30 p <0.0001, Nic 0.15 vs. Nic 0.30 p = 0.0067, Bonferroni’s post hoc test; Suppl fig 1 B2). However, nicotine showed no effect on the total number of chains suggesting that the drug does not change the motivation to perform the task (F (2, 52) = 1.834, p = 0.1751; Suppl fig 1 B3). Importantly, these results also show that despite the reduction in general activity, the motivation to perform the task was no different after nicotine injections. Actually, nicotine 0.15 mg/kg decreases the general activity in females while increases their efficacy, suggesting that the general activity does not impact the number of rewards obtained as long as the mice pay attention to the cue. Finally, nicotine did not affect the latency to collect the reward in females, suggesting that nicotine did not modify the sensitivity to the reward and the reward seeking (F (2, 52) = 1.488, p = 0.2361; Suppl fig 1 B4).

These data suggest that a single administration of a low concentration of nicotine (0.15 mg/kg), but not a higher dose, increases optimal performance in the general population of males and females by increasing the proportion of achieved goals (rewarded chains) independently of motivation and seeking for the reward, validating its pro-cognitive effect in the FCNcue task. Moreover, these data allowed us to identify the optimal responses and the impulsive ones as behavioural biomarkers that are specifically related to attentional control and that modified by a single administration of nicotine.

### 3.3. The pro-cognitive effect of nicotine in females is contingent on their innate attentional control

Our results showed that one of the strengths of the FCNcue task is to unmask inter-individual variability in cognitive performance in a population of male and female mice. An intriguing question is to know if the pro-cognitive effect of nicotine is similar among same-sex individuals with different attentional profiles. We therefore split male and female groups in three sub-populations: impulsive, medium, and good performers, based on their optimal responses in the FCNcue test in control conditions (FCN6cue performances after saline injections, see Material and Methods).

The significant correlations of the percentage of optimal *versus* premature responses show the continuum of attentional performances that male and female mice display under the FCNcue test (Pearson correlation coefficient, for males: r = −0.7521, p < 0.0001; for females r = −0.7940, p < 0.0001; Fig. 4 A1 and Fig 4 B1, respectively). However, this correlation disappears in the group of males after the injection of nicotine 0.15 mg/kg (Pearson correlation coefficient, for males: r = −0.4052, p = 0.0863, Fig 4 A2) whereas the group of females under nicotine 0.15 mg/kg influence keeps showing a significant continuum of responses (Pearson correlation coefficient, r = −0.8871, p < 0.0001) suggesting that the pro-cognitive effect of nicotine 0.15 mg/kg is not ubiquitous. These results suggests that in males, nicotine improved the performance in all subjects, independently from their endogenous attentional performance in control condition. However, in the group of females, even though some of them benefit from nicotine pro-cognitive effect, others do not. In the figures 4 A3 and B3 we can see more clearly that even though nicotine improves attention in males and females as a group (Paired t test, p = 0.0002 and p = 0.0006, respectively) the pro-cognitive effect of nicotine in the group of females differs depending on their basal attentional performance. Thus, medium and good performer females improve their attentional control from 36.7 ± 13.4 % to 53.9 % optimal responses (p = 0.0003, Paired t test); however, impulsive females do not seem to beneficiate from nicotine pro-cognitive effect (p = 0.9822, Paired t test; Fig 4 B3). On the contrary, impulsive males benefit from nicotine pro-cognitive effect (optimal responses in control 19.7 ± 9.9 % to 53.8 ± 15.3 %, p = 0.0001, Paired t test; Fig 4 A3).

**Figure 4.**
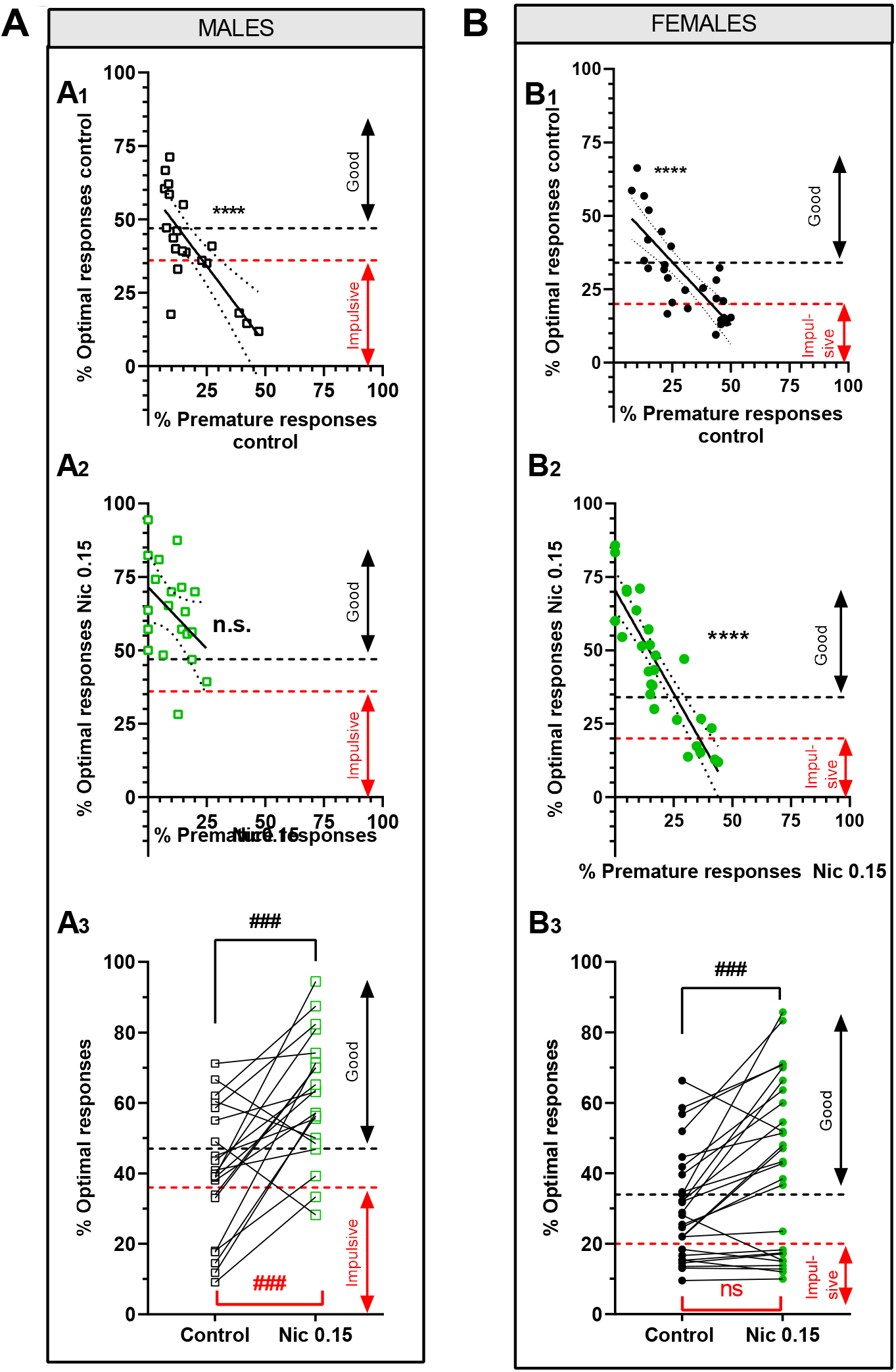
The pro-cognitive effect of nicotine varies according to gender and attentional control profile. To better explore interindividual differences in the pro-cognitive effect of nicotine the populations of males and females were divided in three groups regarding their optimal responses in control conditions: impulsive (lower thirtieth percentile), medium and good performers (higher thirtieth percentile). In these graphs the subpopulations of interest regarding their attentional profile, impulsive and good performers, are those under the red and above the black discontinuous lines, respectively. A1-B1) Correlation between optimal and premature responses shows a continuum of attentional profiles in male and female mice in control conditions: from impulsivity to good attentional control (Pearson correlation coefficient, males r = −0.7521, females r = −0.7940). A2) Nicotine 0.15 mg/kg improves the attentional control in general in males, which makes that all performances improve and there is no correlation between optimal and premature responses (Pearson correlation coefficient r = −0.04052). B2) After nicotine 0.15 mg/kg there is still a significant correlation between optimal and premature responses in females (Pearson correlation coefficient r = −0.08871). Note that the continuum of responses in this group is still present (significant correlation). A3-B3) Percentage of optimal responses in control conditions and after nicotine 0.15 mg/kg injection in the FCNcue test. Male (A3) and female (B3) groups show a significant increase of optimal responses after nicotine injection. However, if we look at the subpopulations in control conditions, nicotine seems to increase the rate of optimal performances in most males (note most connecting lines go up), including the most impulsive ones (red arrow, red statistics) but not the ones of impulsive females (red arrow, red statistics) (**** p < 0.005 one sample t test, linear regression different from 0); ### p < 0.005 Paired t test).

These results suggest that the pro-cognitive effect of nicotine is not ubiquitous: it depends on the sex and, in the case of females, is contingent on their innate attentional control. The implications that this heterogeneity of nicotine pro-cognitive effect has in the vulnerability to develop a preference for nicotine in males and females is shown in the next section.

### 3.4. Nicotine induces CPP in a larger population of males than of females

The conditioned place preference (CPP) procedure is commonly used to study drug reward in rodents, an important process in the initiation of drug addiction. In the CPP experiment, the same animals that underwent the FCNcue test received nicotine in one environment or ‘context’ and saline in another environment. In the CPP we injected nicotine 0.30 mg/kg since we had previously shown that it induces preference for nicotine in mice (16).

The results from the nicotine-induced conditioned place preference (CPP) in males and females are depicted in figure 5 A1 and 5 B1, respectively. Wilcoxon analysis comparing test to pretest total time showed that conditioning with nicotine increased the time spent in the drug-paired compartment during the test in both, the group of males and the group of females (p< 0.0001, Wilcoxon paired comparison, for both, males and females).

**Figure 5.**
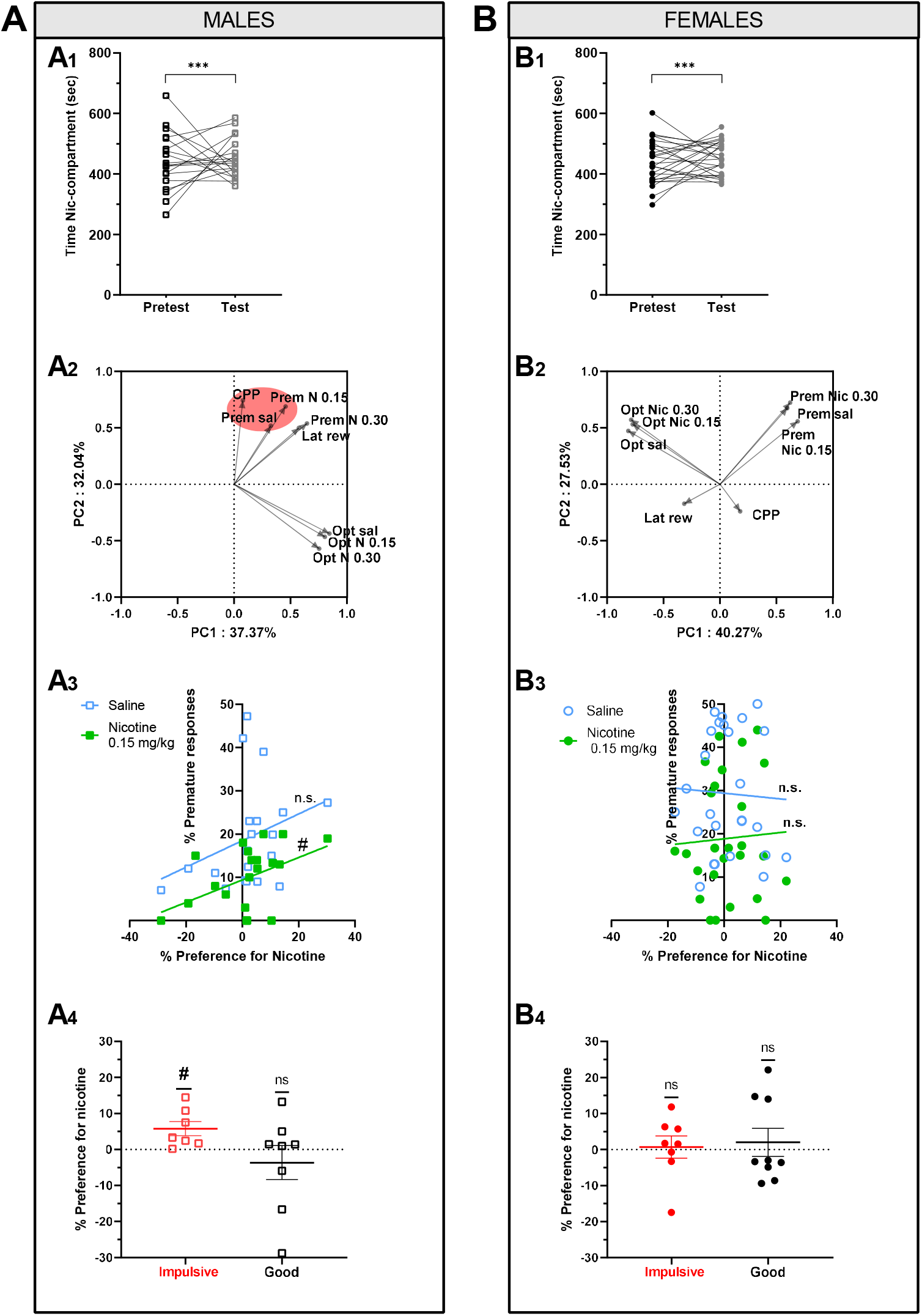
Attentional control profile and pro-cognitive effect of nicotine predict vulnerability to developing nicotine addiction in males but not in females. Multidimensional analysis taking into account behavioural parameters of FCNcue and CPP performances were performed to define behavioural biomarkers that may predict vulnerability to nicotine addiction. A) Multidimensional analysis taking into account behavioural parameters of FCNcue and CPP performances in males. A1) Comparison of the time that the mice spent in the nicotine-paired compartment in the pre-test and test sessions. Conditioning with nicotine generally increased the time spent in the drug-paired compartment during the test. Note that at the individual level, some males show preference (connecting lines going up) while others show aversion (connecting lines going down). A2) Principal Component Analysis (PCA) taking into account behavioural parameters obtained from the FCNcue test (Opt sal= % optimal responses in control conditions, Opt Nic 0.15 = % optimal responses after nicotine 0.15 mg/kg injection, Opt Nic 0.30 = % optimal responses after nicotine 0.30 mg/kg injection; Prem sal,prem Nic 0.15 and Prem Nic 0.30 = % premature (impulsive) responses in control, after nicotine 0.15 and 0.30 mg/kg injection, respectively; Lat rew = latency to the reward in control conditions) and the CPP for nicotine (CPP = % preference for nicotine showed in the CPP test). Note that the PCA showed segregation of preference for nicotine and premature responses (red area), suggesting that in males, preference for nicotine in the CPP is associated with impulsive behaviour in the FCNcue test. A3) Male mice showed a positive correlation between the rate of impulsive responses in the FCNcue test and the preference for nicotine as measured by the CPP when they were under the influence of nicotine 0.15 mg/kg and not in control. A4) When we analysed the preference for nicotine (CPP) in the different subgroups of mice regarding their attentional performance profile, only the subgroup of impulsive males, but not good performers displayed significant preference for nicotine. B) Multidimensional analysis of FCNcue and CPP performances in females. B1) Comparison between test and pre-test showed that conditioning with nicotine increased the time spent in the drug-paired compartment during the test. B2) Principal component analysis showed no segregation of CPP and FCNcue test results, suggesting that in females, attentional performances are not related to nicotine rewarding properties. B3) Females showed no correlation between the preference for nicotine and the rate of impulsive responses in the FCNcue test neither in control conditions, not under the influence of nicotine. These results show that nicotine pro-cognitive effect do not have any impact on the development of preference for nicotine in the CPP. B4) When we analysed the preference for nicotine (CPP) in impulsive and good performer female mice we could not find any significant different between subgroups. Lines and errors represent mean ± s.e.m. *** p < 0.005, Wilcoxon paired test; # p < 0.05, one sample t test (different from “0”).

When we compared the proportion of males and females that showed a preference for nicotine, results show that there is heterogeneity in the reward properties to nicotine between sexes: 76% males show preference for nicotine while in the case of females only 41% show preference for the drug (p = 0.0199, Fisher’s exact test).

### 3.5. Attentional control profile and pro-cognitive effect of nicotine predict nicotine rewarding properties in males but not in females

In an effort to analyse together the main cognitive features that defined attentional control in the FCNcue test and preference for nicotine in the CPP, we performed a multidimensional analysis. To do this, we applied principal component analysis (PCA) as an unbiased statistical analysis to summarize the data variations. Our PCA analysis was based on eight behavioural parameters (all continuous) reflecting the intrinsic properties of the 21 males and the 27 females included in the study. These parameters were the percentage of optimal and premature responses in control conditions and after nicotine 0.15 and 0.30 mg/kg, the latency to the reward, and the score of preference for nicotine. In the PCA analysis for the group of males, the first principal component (PC1) is the major variance, accounting for 37.37 % of the overall variation of our observations as compared to 32.04 % in PC2 component. Interestingly, PCA showed segregation of premature responses and CPP (Fig 5 A2), suggesting that in males, preference for nicotine in the CPP is correlated to impulsive behaviour in the FCNcue test. In the case of females, PC1 accounted for 40.27% of the overall variation of our observations as compared to 27.53 % in PC2 component. The multidimensional analysis showed that there is no interaction between any of the FCNcue parameters and preference for nicotine in females (Fig 5 B2).

Results obtained from the PCA together with the differences in the pro-cognitive effect of nicotine between impulsive males and impulsive females (Fig 4 A3 and 4 B3) made us to focus on premature, i.e. impulsive, responses as a behavioural biomarker related to attentional control, pro-cognitive effect of nicotine and sensitivity to nicotine reward. Therefore, we analysed directly the correlation between the impulsive responses in the FCNcue test and the preference (or aversion) for nicotine showed in the CPP. In control conditions, there was no significant correlation between impulsive responses and preference for nicotine in males or females (Pearson correlation coefficient for males r = 0.3327, p = 0.1406; for females r = −0.0451, p = 0.8267; Figs 6 A3 and 6 B3, blue). However, after nicotine 0.15 mg/kg injection, we found a positive significant correlation between the percentage of impulsive responses and the preference for nicotine in males (Pearson correlation coefficient r = 0.5404, p = 0.0400; Fig 5 A3, green) but not in females (r = 0.0496, p = 0.8090; Fig 5 B3, green). These results suggest that those males that are more impulsive in the FCNcue test, which benefit the most from the pro-cognitive effect of nicotine (Fig 5 A3, upper values in the premature responses axis, in blue, go down after nicotine, in green), might develop a positive affective memory associated with nicotine that induces a larger preference for the drug in the CCP. However, impulsive females, which do not benefit from nicotine pro-cognitive effect, do not show any link between attentional control, nicotine pro-cognitive effect and preference for nicotine. Finally, we analysed if the preference for nicotine in the subpopulations of impulsive (poor performance) and good performers was different. Remarkably, all impulsive males, but not the good performers, showed a significant preference for nicotine (p = 0.0280 for impulsive and p = 0.4619 for good performers, One sample t test; Fig 5 A4). In females, neither the impulsive or good performer subgroups showed a clear preference for nicotine (p = 0.8263 and p = 0.6207, respectively, One paired t test; Fig 5 B4). These results suggest that in a population of male mice, those showing naturally poor inhibitory control, i.e. impulsive, are more susceptible to develop nicotine addiction. Whereas in the case of females, there is no link between the attentional profiles and nicotine rewarding properties.

## Discussion

We sought to examine whether the effects of nicotine on cognitive performance in mice could be predictive of sensitivity to nicotine reward and, if so, whether this characteristic was gender dependent. To do this, we adapted the FCNcue test to study attention and inhibitory control capacities in male and female mice. We were able to observe both inter-individual and sex differences in attentional processes. A single injection of a low dose of nicotine improved attentional performance in the male population, independently of their attentional control profile. However, in females, the same dose did not improve performance in animals who had poor baseline attentional control, namely impulsive. Moreover, multidimensional analysis of parameters obtained from the FCNcue and CPP tasks showed that healthy males with poor basal attentional control profiles were more responsive to the rewarding properties of nicotine and may therefore be more vulnerable to developing a nicotine addiction than males showing good attentional control. Females, on the contrary, show a completely different behavioural pattern. Specifically, the preference for nicotine does not depend at all on the attentional parameters measured. These results may open new avenues to study sex-specific physiological and neurobiological mechanisms of vulnerability to nicotine addiction.

We had previously developed the “cued fixed consecutive number” task (FCNcue), which enables the characterization of attention and inhibitory control, for rats (13,14). The principle of the FCN is simple and efficient: it measures the ability of subjects to perform a chain of sequential actions to obtain a reward. The addition of a cue (FCNcue) light to the original FCN task (18) is central to the aim of dissociating attention from the behavioural inhibition deficit that gives rise to impulsivity (13). With this modification to the task, the animal does not rely on internal signals such as timing (18) for successful performance of the chain of action, but rather on its attention to cue detection to start and to the cue light turning off to switch to the next step. Here, we adapted the FCNcue test to mice and examined the pro-cognitive effect of nicotine on male and female mice. The variability we observed in the parameters related to baseline attentional performances of male and female mice in the FCNcue task (optimal and premature responses, efficiency) and the subsequent improvement following injection of pro-cognitive drugs are both consistent with previous studies performed with rats (13,18–20). Our present results thus constitute a robust proof of concept for the adaptation of the FCNcue task to mice, which opens to others and us new possibilities to study specific neuronal circuits underlying attentional control thanks to the wide range of transgenic mice lines available.

Population grouping on the basis of attentional performance in the FCNcue task makes it ideal for broader application in the investigation of behavioural biomarkers for other clinical conditions characterized by poor attentional control, such as ADHD, or schizophrenia (21). Our study of inter-individual heterogeneity in mice not only contributes to explaining differential susceptibility to developing psychiatric conditions (such as drug addiction), it also highlights methodological problems that can lead to inappropriate or over-simplistic interpretations (22). Studies in rodents allow us to use approaches that are not conceivable in humans; still we ought to develop animal models that are as close as possible to human studies. The reliability showed in our study of attentional performance measurement across littermates and sexes and the effect of stimulants support the utility of the FCNcue as a means by which to assess attention in rodents in a comparable form to the Continuous Performance Test (CPT) used in humans (23,24). Therefore, our results demonstrate that FCNcue has translational validity for investigating attentional deficits in psychiatric disorders and new cognitive therapies.

It has been suggested that nicotine improves general attention in smokers and that the beneficial cognitive effects of nicotine may have implications for the maintenance of tobacco dependence (25). In their meta-analysis, Heishman and colleagues also suggest that performance facilitation might play a role in the rewarding effects of nicotine during the initiation of tobacco dependence performance; however, this hypothesis had not been empirically tested. Our results show that a single injection of nicotine increases attentional performance in male and female mice, as groups. Here, we also show that attention improves without modifying motivation or sensitivity to natural reward, confirming the pro-cognitive effect of nicotine in mice. A deeper analysis of our data showed that, at the individual level, nicotine improves attentional performance in the population of male mice, even in those that show a majority of premature responses in the control condition, i.e. the impulsive males. However, nicotine does not improve attentional control in impulsive females. These results reveal that the pro-cognitive effects of nicotine are sex-specific. Our results in impulsive females indicate the importance of 1) including both sexes in preclinical studies and 2) analysing animals according to their behavioural phenotype. At the behavioural level, we observe impulsivity in both males and female. However, when we add a neuromodulator (nicotine) their cognitive abilities are not affected in the same manner. This suggests that the source of impulsivity showed in the FCNcue test does not have the same cognitive and/or neurophysiological foundations in males and females. There remains a surprising lack of consensus regarding how sex influences impulsive choice (26,27). Some have related these differences to hormones; however, their results, especially in females are inconclusive (28).

To understand the potential link between the pro-cognitive effect of nicotine and the risk of developing nicotine addiction, we exposed mice that had undergone the FCNcue task to a nicotine-induced conditioned place preference paradigm (CPP). CPP is a widely used protocol for measuring drug abuse liability in humans as well as animals (29). Drug abusers report strong associations with environments in which they use drugs and the acquired incentive properties of drug-associated places can elicit craving and/or relapse in the abstinent addict (30). In line with previous studies, our results in the CPP confirm that males are more sensitive than females to the rewarding properties of nicotine (31). In addition, we show that innate impulsivity, predicts sensitivity to nicotine rewarding properties in males but not in females (32). We also found a discrepancy between the nicotine pro-cognitive dose (0.15 mg/kg) and the dose usually used in the CPP (0.30 mg/kg). We have previously shown that nicotine 0.30 mg/kg induces place preference in mice (16). However, our results in the FCNcue test are clear: nicotine 0.30 mg/kg does not improve attentional control in males or in females. Therefore, we report that in our experimental conditions nicotine’s effective dose to enhance attentional control is lower than that to induce rewarding/ aversive effects. This discrepancy in the pro-cognitive and rewarding effect of the drug has been reported previously for other compounds such as amphetamine, which pro-cognitive optimal dose is 0.30 mg/kg and the dose used in the CPP is usually 2 mg/kg (17,33).

Beyond impulsivity and nicotine addiction, our findings also show that the pro-cognitive effect of nicotine in impulsive males (but not females), leads to a greater sensitivity to nicotine reward in this subpopulation as demonstrated using the CPP task. These results suggest that impulsive males that benefit from a single injection of nicotine in the FCNcue test, would keep a reminiscence memory of the positive effect of the drug (better attentional control = more rewards), predisposing them to “like” the effect of nicotine in the CPP and therefore to develop nicotine addiction. It has been previously suggested that the pro-cognitive effect of nicotine may be linked to tobacco abuse in psychiatric disorders related to an impairment of inhibitory control such as schizophrenia and ADHD (34,35). However, there is still no consensus about the origin of nicotine abuse in these patients: if it acts a self-medication due to its pro-cognitive properties or if they are more sensitive to the rewarding effect of the drug, or both. To our knowledge, this is the first time that pro-cognitive effects of nicotine are linked to nicotine rewarding properties in subjects who naturally show impulsive behaviours. Opening the possibility to use the FCNcue test as a behavioural task to measure natural impulsivity as a biomarker to develop other psychiatric syndromes such as ADHD or schizophrenia. Our data also show that the pro-cognitive effect of nicotine does not seem to have any impact on the vulnerability to develop nicotine addiction in impulsive females. These results would be in the same line as others suggesting that women have a higher sensitivity to the non-pharmacologic effects of nicotine than men (36,37).

To conclude, this study highlights the need to develop preclinical models that consider the inter-individual heterogeneity of behavioural responses and both sexes. Here, we propose that impulsive males, but not impulsive females, benefit from the pro-cognitive of nicotine, which makes them more vulnerable to like the effects of the drug and therefore to seek for it and develop addiction. Overall, this suggests that the cognitive strategies involved in innate impulsivity may not the same in males and females, and the neuronal bases are likely to differ. Now our objective is to investigate by the use of chemogenetics whether specific prefrontal cholinergic circuits are equally involved in attention and inhibitory control in males and females. Such studies are vital in order to disentangle firstly, the underlying biology driving these sex differences in natural impulsivity and secondly, the role of impulsive behaviour in the vulnerability to develop other psychiatric disorders in males and females.

## Acknowledgements

The authors thank Yoan Salafranque for his technical help and Shauna Parkes for her comments on the manuscript.

## Authors’ contribution

MCM, FD, MC and SC developed the concept and design of the study. MCM and FD performed the experiments. MCM, FD and SC participated in data analysis and interpretation of the findings. MCM and SC wrote the paper. All authors critically reviewed content and approved the final version for publication.

---

**Supplementary figure 1.**
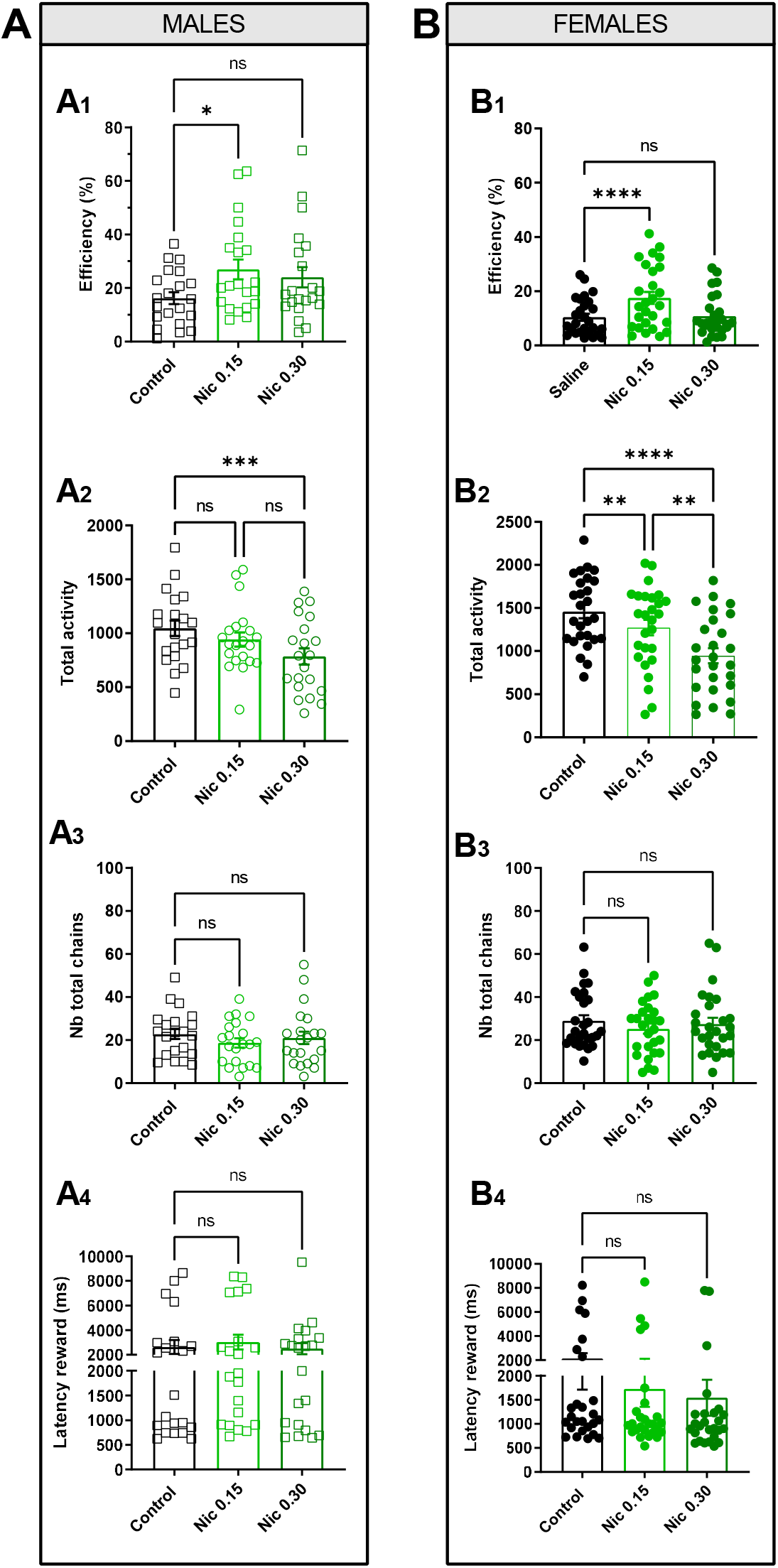
The pro-cognitive effect of nicotine in the FCNcue test is independent of motivation and reward seeking. A) Nicotine effect on efficiency, motivation and reward seeking during the FCNcue test in the group of males. Nicotine 0,15 mg/kg (but not nicotine 0,30 mg/kg) increases efficacy in the FCNcue test in males (A1). Nicotine 0.30 mg/ kg, but not nicotine 0.15 mg/kg, decreases general activity during the FCNcue test (A2). Nicotine does not show any effect on the total number of chains of nose-pokes (motivation to work) (A3) or in the latency to get the reward (-reward seeking) (A4) in males. B) Nicotine effect on efficiency, motivation and reward seeking during the FCNcue test in the group of females. Nicotine 0,15 mg/kg (but not nicotine 0,30 mg/kg) increases efficacy in the FCNcue test in males (B1). Nicotine decreases general activity during the FCNcue test in a dose-dependent way (B2). Nicotine does not show any effect on the total number of chains of nose-pokes (motivation to work) (B3) or in the latency to get the reward (reward seeking) (B4) in females. Note that the decrease in the general activity does not have an impact on the motivation for the task. Bars and errors represent mean ± s.e.m. * p < 0.05, ** p < <0.01, *** p < 0.005, **** p < 0.0001, Bonferroni’s post hoc test.

